## Supplemental material for "Structure and function of *Plasmodium* actin II in the parasite mosquito stages"

### Supplemental tables

**S1 Table.** Data collection and refinement statistics.

| **Data collection** | **Actin II** | **Actin II-JAS** |
|---|---|---|
| Magnification | 75000x | 75000x |
| Defocus range (mm) -Voltage (kV) | -300 | -300 |
| Microscope | Titan Krios | Titan Krios |
| Detector | Falcon 3 | Falcon 3 |
| No. of frames | 46 | 46 |
| Pixel size (Å/pixel) | 1.09 | 1.09 |
| No. of micrographs | 1058 | 3977 |
| **Reconstruction (Relion 3.1beta)** | | |
| No. of helical segments | 47 197 | 272 310 |
| Box size (px) | 328 | 328 |
| Rise (Å) | 28.34 | 28.37 |
| Azimuthal rotation (°) | -166.9 | -166.9 |
| Average resolution (Å) (FSC=0.143) | 3.5 | 3.3 |
| Model resolution (Å) (FSC=0.5) | 3.4 | 3.2 |
| Map sharpening B-factor (Å$^2$) | -111 | -75 |
| **Model building (Phenix 1-19.2-4158)** | | |
| No. of chains | 6 | 6 |
| No. of atoms | 18102 | 18324 |
| No. of amino acid residues | 2232 | 2232 |
| No. of ligand atoms ($Mg^{2+}$, ADP, JAS) | 12 | 18 |
| Average B-factor (Å$^2$) | 49.25 | 38.63 |
| Average B-factor for ligand atoms (Å$^2$) | 37.50 | 46.79 |
| R.m.s.d. bond lengths (Å) | 0.002 | 0.002 |
| R.m.s.d. bond angles (°) | 0.543 | 0.558 |
| CC volume | 0.83 | 0.82 |
| CC masked | 0.85 | 0.84 |
| Molprobity score | 1.17 | 1.12 |
| Clash score | 3.52 | 3.32 |
| Ramachandran plot favored/allowed/outliers (%) | 97.87/2.13/0 | 98.09/1.91/0 |
| Rama-Z (whole/helix/sheet/loop) | 0.45/1.43/0.18/0.65 | 0.64/1.24/0.76/0.43 |
| **Deposition codes** | | |
| PDB | XXXX | YYYY |
| EMDB | EMD-10588 | EMD-10589 |

**S3 Table.** Relative kinetic parameters of actin II polymerization. All values are reported as mean ± standard deviation (actin II seeds: n=3 NP, n= 5 in SS). * Relative $k_-$ calculated using the steady state $C_c$ and relative elongation constant ($k_+$) from slope 2 of nucleated polymerization assays.

| | | |
|---|---|---|
| **Nucleated polymerization (NP)** | | |
| | **Actin II seeds** | |
| | *Slope 1* | *Slope 2* |
| $k_+$ ($s^{-1}$) | 0.04 ± 0.01 | 0.01 ± 0.002 |
| $C_c$ ($\mu$M) | 1.12 ± 0.29 | 0.36 ± 0.19 |
| $k_-$ ($s^{-1}$) | -0.04 ± 0.02 | -0.004 ± 0.003 |
| **Steady state (SS)** | | |
| $C_c$ ($\mu$M) | 0.11 ± 0.02 | |
| $k_-$ ($s^{-1}$)* | -0.003 ± 2e-004 | |

### Supplemental movie legends

**S1 Movie.** Evolutionary conservation of *P. falciparum* actin II in comparison with A actin I from *Plasmodium* spp. The actin surface is colored according to conservation scores; high (purple) to low (variable). Amino acid conservation was estimated using ConSurf server [74].

**S2 Movie.** Twistedness motion of G-actin II and F-actin II protomers. G-actin II (6I4M) structure and our F-actin II model were used to generate the motion using Morph conformation in Chimera [68]. There is no crystal structure of actin II available, G-actin from *P. berghei* shares 92.6% sequence identity with actin II. The movement of residues around the nucleotide-binding site is shown.

**S3 Movie.** The twist angles of the mass centers of SDs ($\theta$) of F-actin II protomers. Superimposed structures of JAS-stabilized F-actin II (medium purple) with **(a)** G-actin II (PDB: 6I4M) (lemon) and **(b)** actin I in F form (6TU4) (light blue). The spheres represent the center mass of each subdomain. The center of mass was calculated in Pymol v1.7.4 and visualized in Chimera [68]. Structures are aligned using SD3. F, filament, and G, globular.

**Supplemental figure legends**

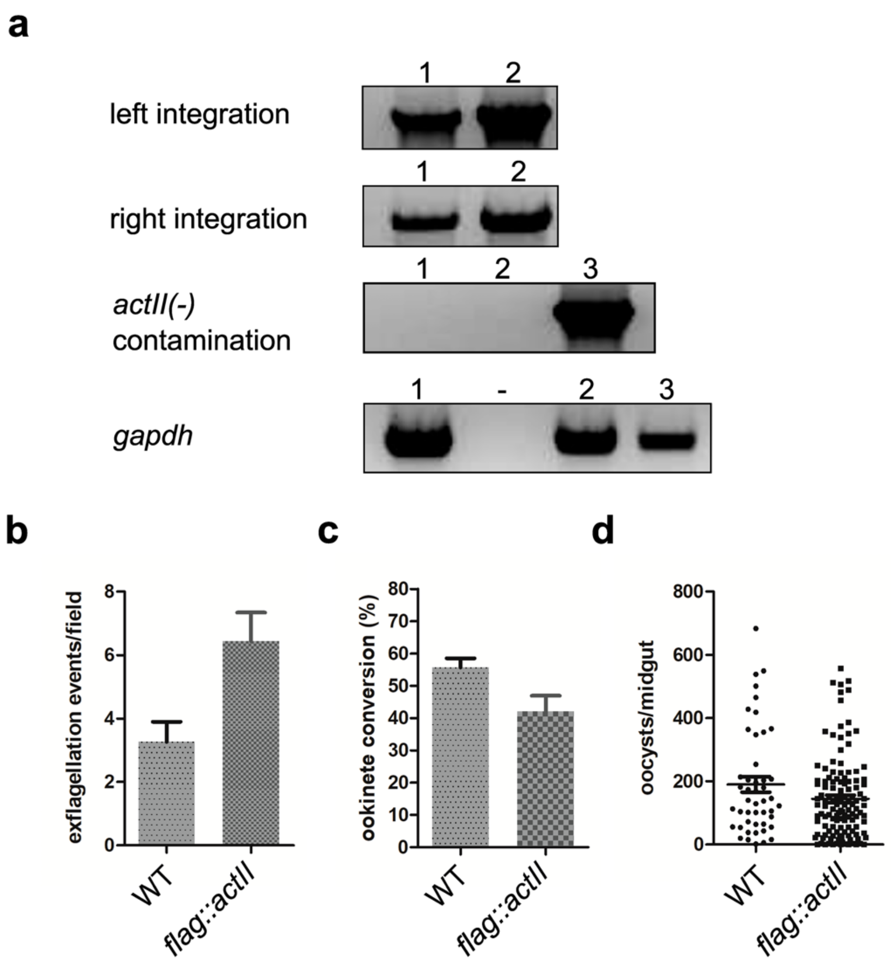


**S1 Fig. (a)** Genotyping *flag-actII* cloned line. Left integration of the construct was verified with primer pair A2F2 and A2R and right integration with the primer pair DHFR and mCherryR. To control for absence of the *actII(-)* parasites the primer pair A2F1 and mCherryR was used. Lane 1 and 2: *flag::actII,* lane 1, 135 ng, lane 2, 50 ng template; lane 3: *actII(-)*. Quality control of gDNA used the *gapdh* primer pair of the same samples. **(b)**. **(b-e)** Phenoptyic analysis of *flag::actII* compared to WT. **(b)** Exflagellation analysis; average of three experiments of the WT and four of the mutant. **(c)** Ookinete conversion; three experiments of each strain. Error bars in **(b)** and **(c)** are S.E.M. **(d)** Number of oocysts/midgut. The oocysts were stained with Cap380 antibody and all oocysts irrespective of size are plotted. Differences in **(b-d)** are not significant, Student's t-test for **(b)** and **(c)**, Mann-Whitney for **(d)** and **(e)**.

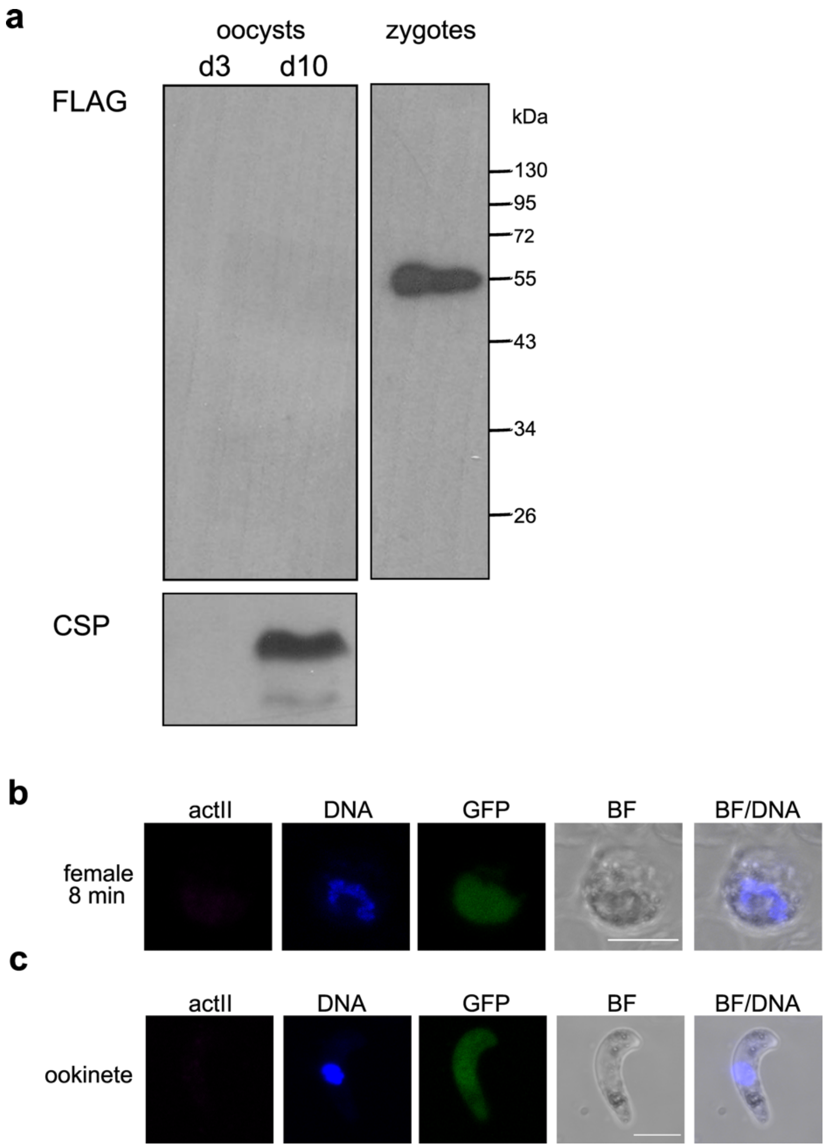


**S2 Fig. (a)** Western blot of extracts from midguts infected with the line expressing FLAG::actinII. Midguts were dissected on day 3 and 10 post blood feeding and crude extracts of 20 and 13 midguts, respectively, were loaded in each lane. Right panel is a positive control with zygote extracts. The samples were run on the same gel and the blot was probed with the anti-FLAG antibody. A duplicate blot was probed with anti-CSP antibody as a loading control; it only gave a signal for the day 10 sample. **(b,c)** Immunolabeling of *flag::actII* female gamete 8 min p.a. **(b)** and ookinete **(c)**. No signal was detected with the anti-FLAG antibody. The background GFP signal (green) is constitutively expressed in this line. DNA was stained with Hoechst 33342 (blue). Scale bars, 5 µm. The female gamete originates from the same experiment as the 8 min sample of male gametes in **Fig 1** (lower panels).

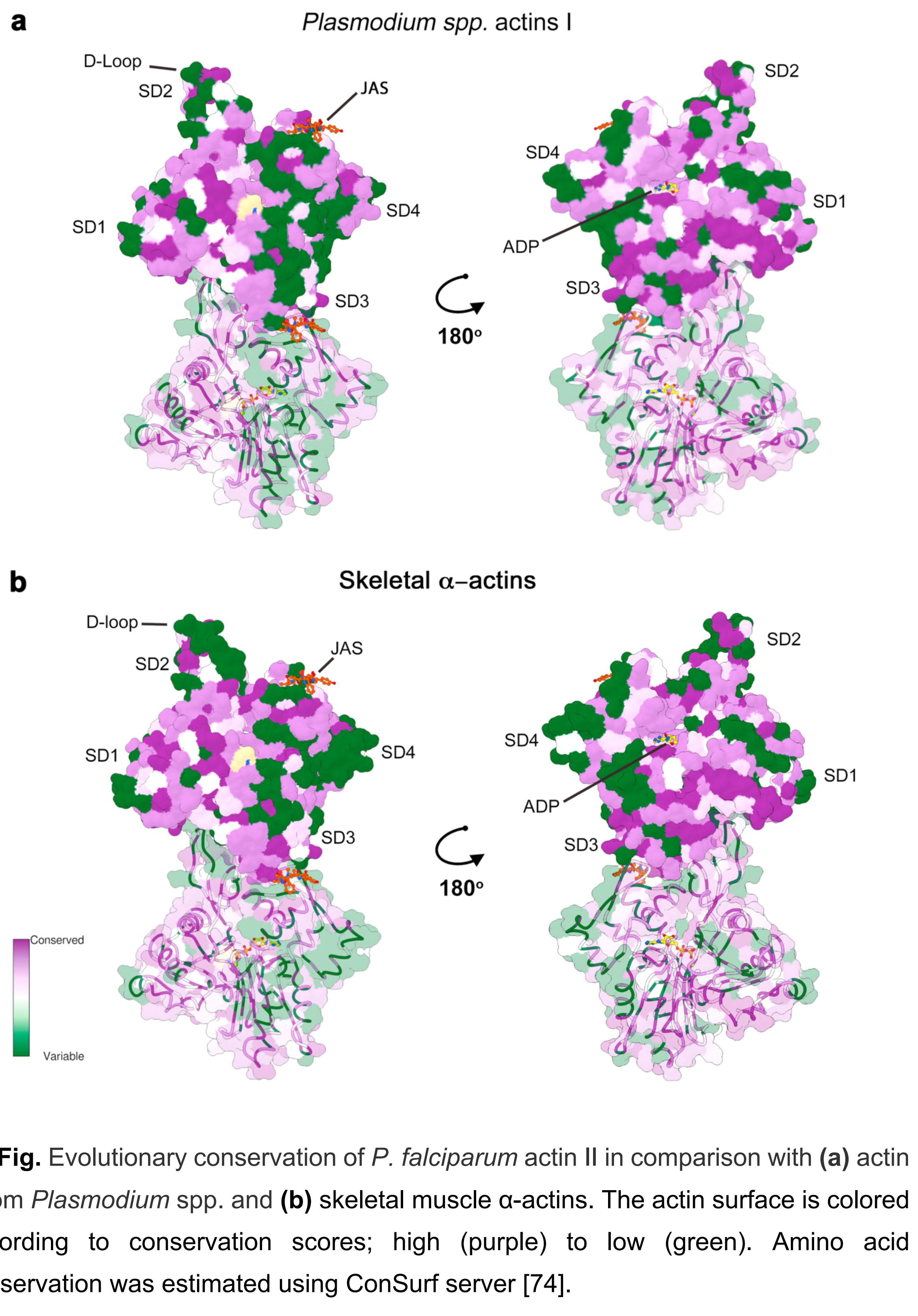


**S3 Fig.** Evolutionary conservation of *P. falciparum* actin II in comparison with **(a)** actin I from *Plasmodium* spp. and **(b)** skeletal muscle α-actins. The actin surface is colored according to conservation scores; high (purple) to low (green). Amino acid conservation was estimated using ConSurf server [74].

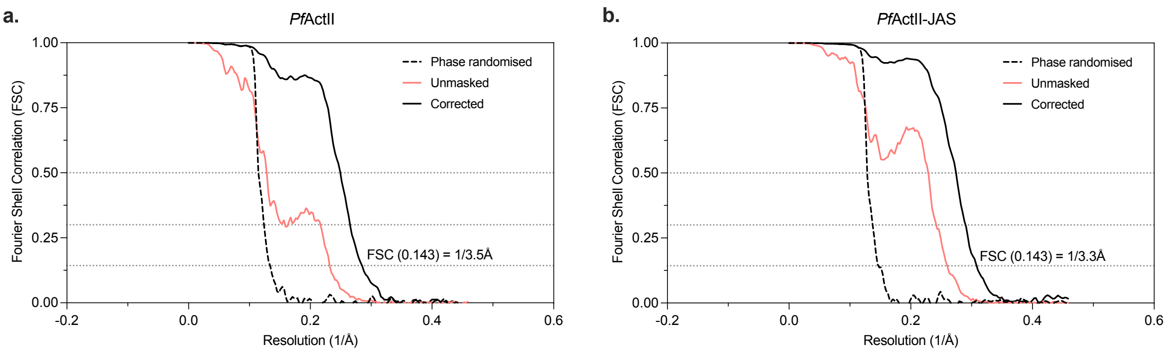


**S4 Fig.** Fourier shell correlation plot of actin II and JAS-stabilized actin II. The corrected curve was calculated from independently refined half-datasets with a soft mask filtered to 15 Å in Relion [75].

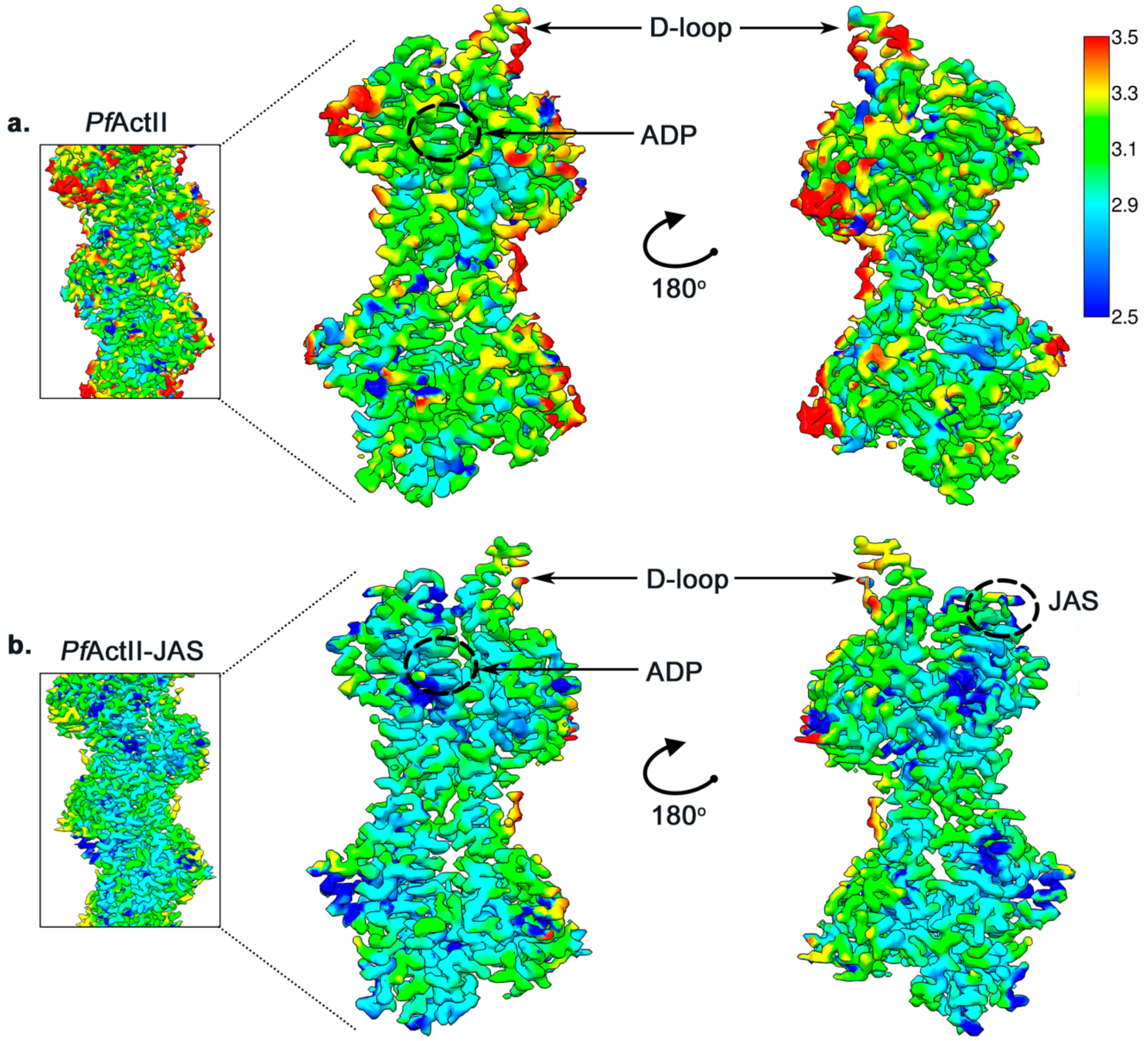


**S5 Fig.** Local resolution of actin II **(a)** and JAS-stabilized actin II **(b)**. The local resolution estimation is based on Fourier shell correlation threshold 0.143 calculated with Blocres in the Bsoft software package, applied to the final sharpened map. The left panel shows a central section of the filament. On the right, two adjacent protomers are shown and, the ligands densities are highlighted [66].

| Aggregation state | Structure | θ (°) | b2 (Å) | d2–4 (Å) | d3-4 (Å) | c (Å) |
|---|---|---|---|---|---|---|
| | *Pf*ActII | 10.1 | 5.4 | 24.8 | 8.8 | 9.0 |
| | *Pf*ActII-JAS | 9.7 | 5.7 | 24.8 | 9.1 | 9.1 |
| Filament | *Oc*Act (5JLF) | 8.2 | 5.8 | 23.1 | 8.0 | 9.1 |
| | *Pf*ActI-JAS (5OGW) | 7.2 | 6.2 | 24.2 | 8.4 | 8.8 |
| | *Pf*ActI-JAS (6TU4) | 6.2 | 5.7 | 24.9 | 9.1 | 8.8 |
| | *Gg*Act (6DJO) | 8.8 | 5.7 | 23.4 | 8.5 | 8.4 |
| | *Gg*Act-JAS-cLys (5OOC) | 8.6 | 5.3 | 23.1 | 8.1 | 8.8 |
| | *Mm*Act (6KLL) | 7.7 | 5.5 | 23.1 | 8.7 | 8.4 |
| | *Oc*Act (1J6Z) | 27 | 5.6 | 26.7 | 8.1 | 11.1 |
| Monomer | *Pf*ActI (6I4E) | 19.5 | 5.1 | 27.6 | 8.3 | 10.3 |
| | *Pb*ActII (6I4M) | 21.4 | 5.5 | 25.9 | 8.2 | 8.9 |
| | *Bt*β-Act (1HLU) | 17.6 | 8.2 | 30.5 | 8.0 | 15.8 |

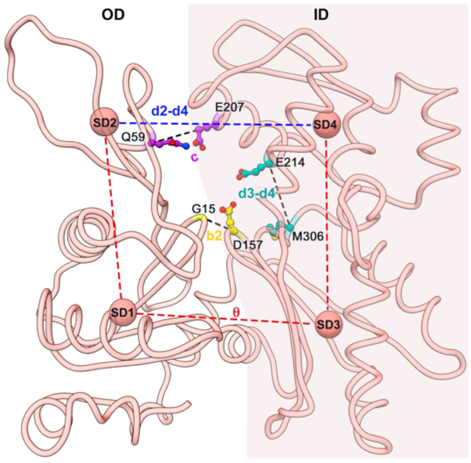


**S6 Fig.** The distance of the twist angles of the centers of mass of SDs ($\theta$), the phosphate clamp distance (b2), the distance of SD1 and SD2 (d2-4), SD3-SD4 (d3-4), and cleft mouth (c) calculated for different actin structures. The dihedral angle of subdomains in the JAS-stabilized actin II model is rotated 11.8° relative to the crystal structure of G-actin II-gelsolin (PDB: 6I4M), 2.4° relative to JAS stabilized F-actin I (5OGW), and 3.4° relative to the JAS stabilized F-actin I in complex with myosin A (6TU4).

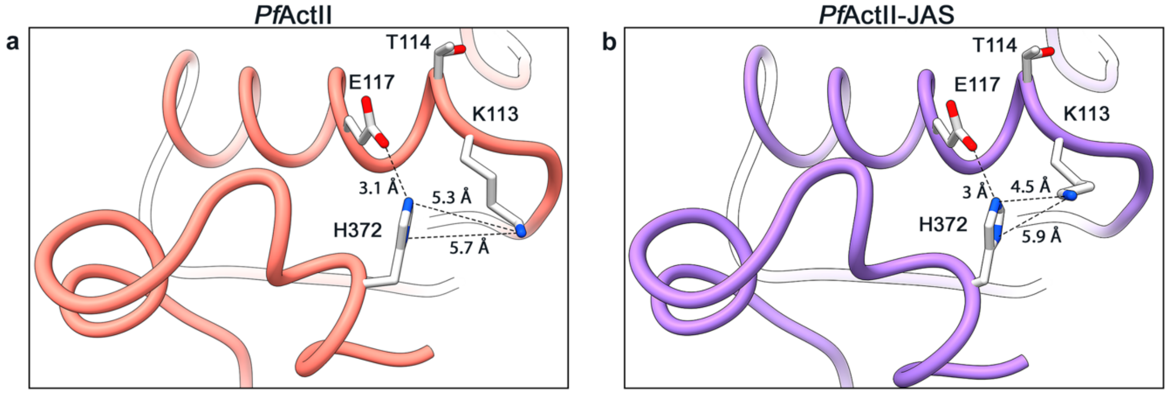


**S7 Fig.** C termini of F-actin II and JAS-stabilized F-actin II. H372 interacts with E117. **(a)** In F-actin II, the ring of H372 is oriented towards the E117. **(b)** In F-actin II, the N1 of the H372 turns towards K113. The distance between H372 and K113 is more than 5 Å in both structures.

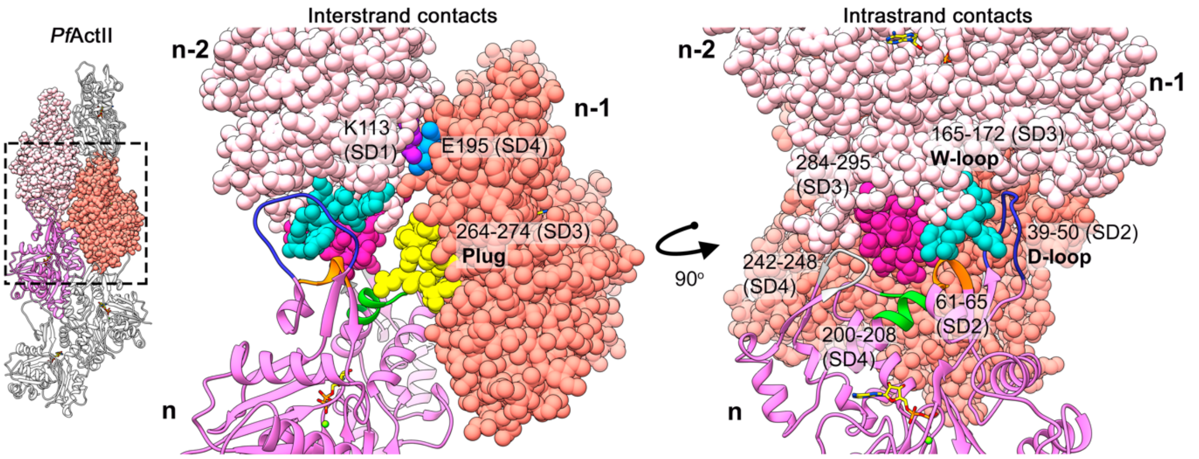


**S8 Fig.** Intrastrand and interstrand interactions of F-actin II and other filamentous actin structures [9,22–24]. Two protomers are represented by spheres n-1 and n-2 (salmon and pink); the silhouette of the third protomer (n) is visualized by ribbon representation in Chimera.

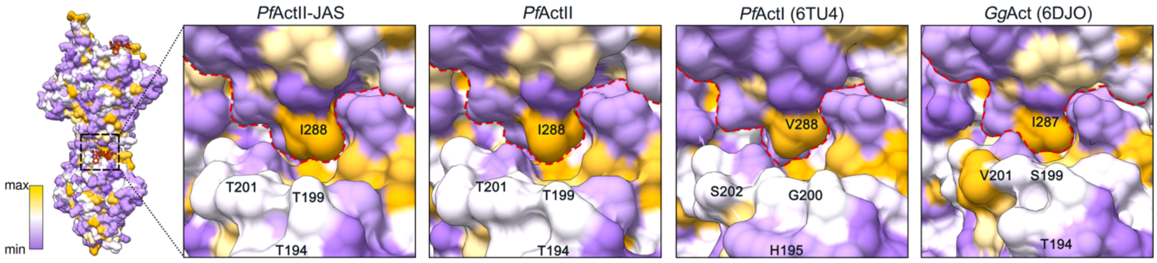


**Fig S9.** Intrastrand contacts near the JAS-binding site. **(a)** *P. falciparum* F-actin II, **(b)** JAS-stabilized F-actin II, **(c)** *P. falciparum* F-actin I (6TU4), **(d)** filamentous skeletal muscle α-actin (6DJO). In actin II, I288 inserts into a groove in the adjacent protomer, resembling a lock-key interaction, like in canonical actins. The actin surface is colored according to hydrophobicity; high (yellow), medium (white), and low (purple).

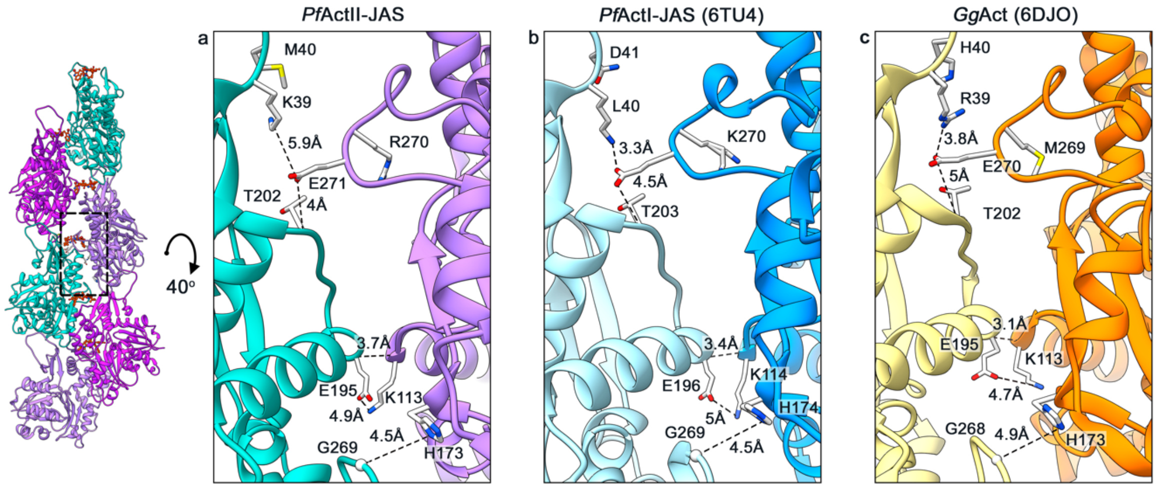


**S10 Fig.** Lateral contacts of protomers from a different strand of **(a)** F-actin II-JAS, **(b)** F-actin I (6TU4), **(c)** *O. cuniculus* F-actin (6DJO). Distances between the residues are indicated in black dashed lines.

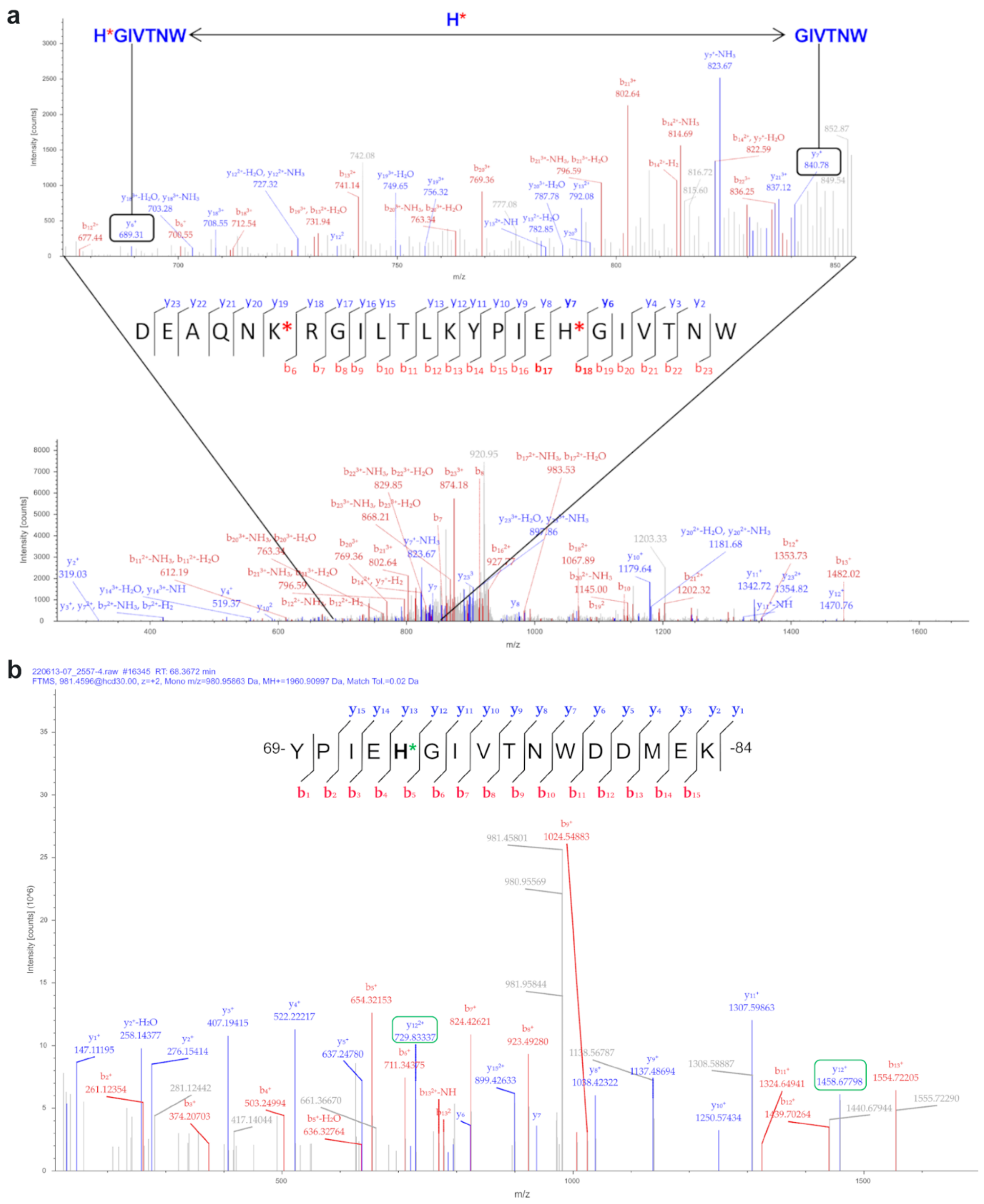


**S11 Fig.** MS/MS spectrum of H73 methylation of *Plasmodium* spp. Actin II **a.** MS/MS spectrum of recombinant actin II. The peptide (56-79) has a methylated group (green star) at 1960.9. Methylated double and triple-charged signals are highlighted in a green box. **b.** MS/MS spectrum of the *Pb*ActII peptide (56-79) obtained after AspN digestion. The peptide sequence is reported, and the red star indicates the presence of a methylation group. b and y series ions detected are reported as red and blue peaks, respectively, in the acquired spectrum such as in the panel containing the peptide sequence. The upper panel shows a zoomed region of the MS/MS spectrum in which y6 and y7 ions are highlighted: the m/z difference between these ions corresponds to a methylated histidine and allows to position of a methylation group on H73 of the actin II.

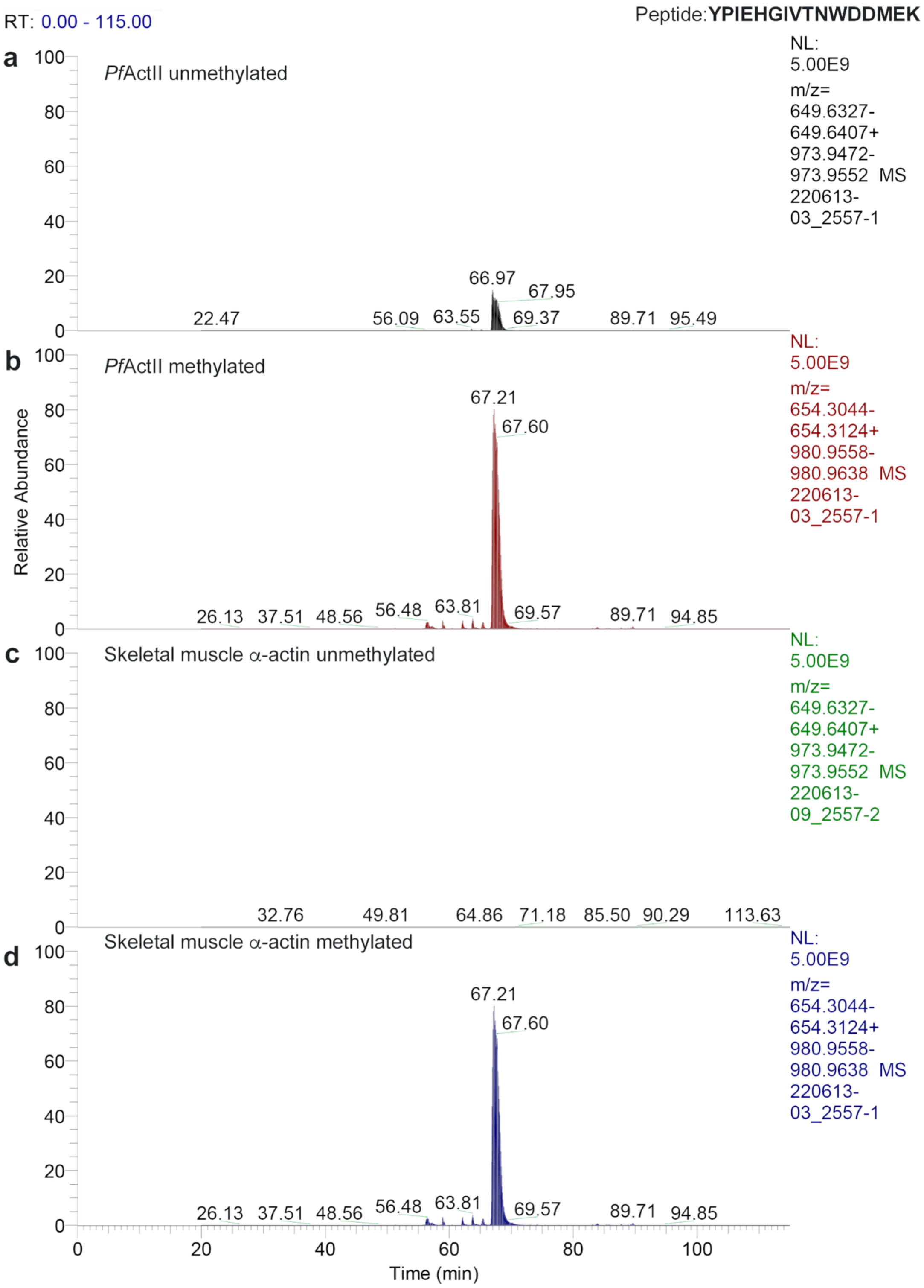


**S12 Fig.** Relative abundance of methylated and unmethylated peptide (56-79) obtained from the recombinant actin II. **a.** Unmethylated and methylated actin II **(b)**. As positive control panels, **c** and **d** show the unmethylated and methylated skeletal muscle α-actin.

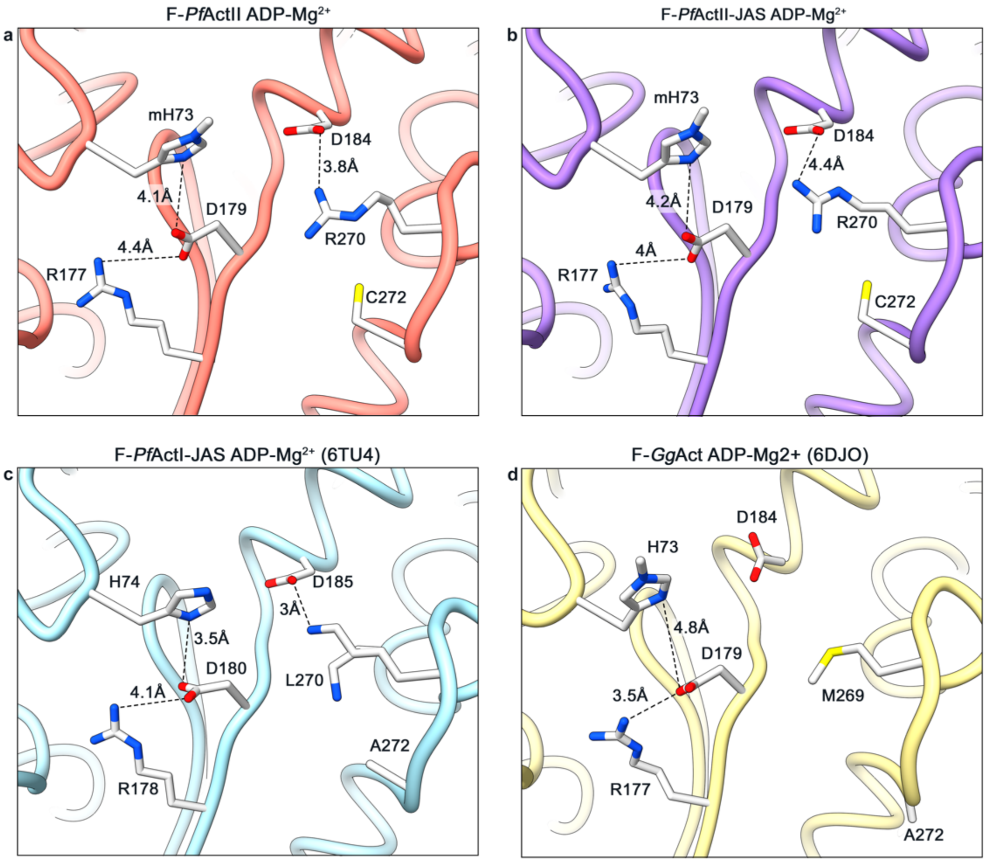


164
165 **S13 Fig.** The orientation of the A-loop in actin II. **(a)** F-actin II, **(b)** F-actin II-JAS, **(c)**
166 F-actin I-JAS (6TU4) and **(d)** *G. gallus* F-actin (6DJO). All structures are in the Mg-
167 ADP state and show a 1b conformation of the A-loop. The most probable ionic and
168 hydrogen bonds are indicated with dashed lines.

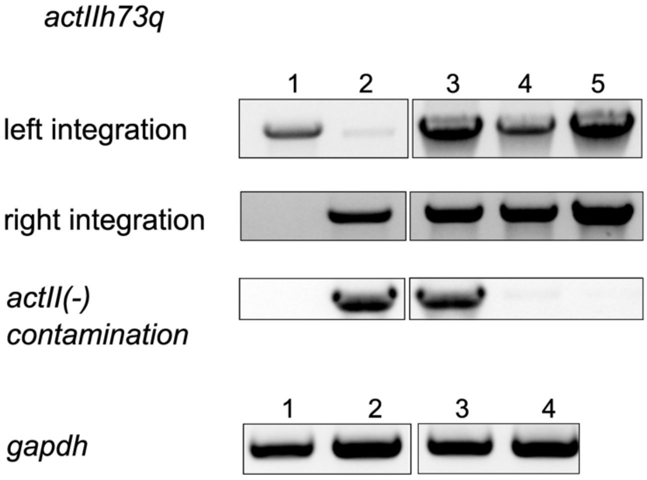


**S14 Fig.** Genotyping of *actIIh73q*. Left integration of the construct was verified with primer pair A2F2 and A2R and right integration with the primer pair DHFR and mCherryR. To control for absence of the actII(-) parasites the primer pair A2F1 and mCherryR was used. **1:** WT; **2:** positive control; **3:** transfected parasites with actIIh73q construct; **4 and 5:** actIIh73q clone 1. For quality control of the gDNA GAPDH primer pair was used.

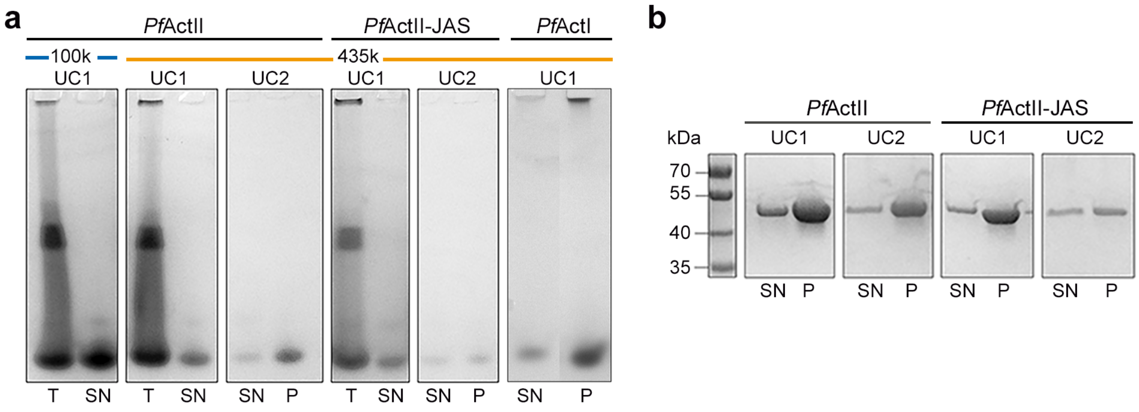


**S15 Fig.** Two steps of sedimentation assay. **(a)** Native PAGE of *Plasmodium* actins. Samples were polymerized overnight, pellet (P) and supernatant (SN) were separated by ultracentrifugation at 4°C for 1 h (UC1). The SN fraction was re-pelleted 16 h after the initial ultracentrifugation (UC2). **(b)** SDS-PAGE of the two steps of ultracentrifugation at 435000 *g* of actin II.

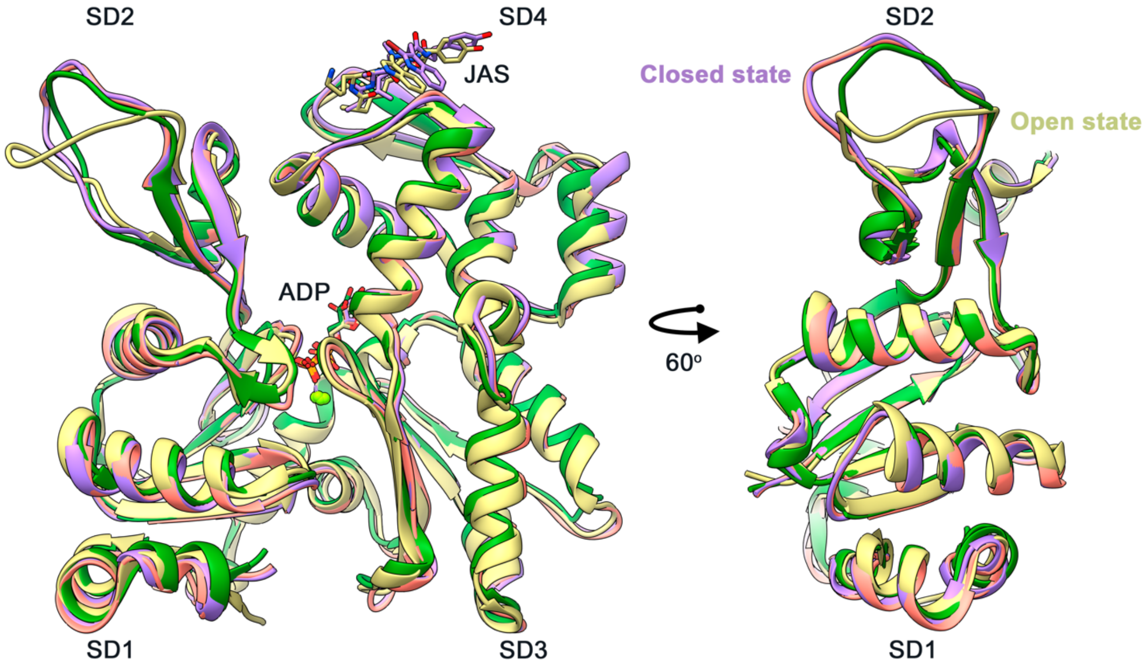


**S16 Fig.** D-loop conformation in actin II. Superimposed structures of JAS-stabilized F-actin II, G-actin II, JAS-stabilized *O. cuniculus* F-actin (5OOC), and *O. cuniculus* F-actin (5ONV). JAS is not altering the conformation of the D-loop of actin II like in actin from *O. cuniculus*, in which the D-loop has an open conformation.

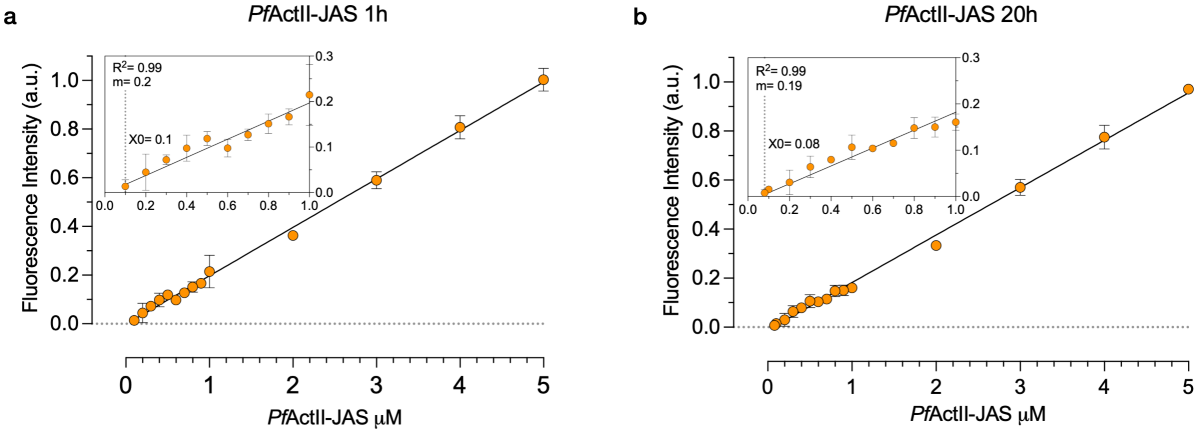


**S17 Fig.** $C_c$ plots of actin II ($n^b$=2). **a.** Samples after 1 h of incubation at 4°C. **b.** Samples after 20 h of incubation at 4°C. The data in both panels fit in a linear regression equation. X0 represents the critical concentration. The lower concentration points are magnified on the left. Error bars represent the standard deviation, a.u.= arbitrary units. $n^b$= biological replicates with 3 technical replicates per experiment.
